# The interaction of phylogeny and community structure: linking clades’ ecological structures and trait evolution

**DOI:** 10.1101/404111

**Authors:** William D. Pearse, Pierre Legendre, Pedro Peres-Neto, T. Jonathan Davies

## Abstract

1

**Aim:** Community phylogenetic studies use information about species’ evolutionary relationships to understand the processes of community ecological assembly. A central premise of the field is that species’ evolution maps onto ecological patterns, and phylogeny reveals something more than species’ traits alone. We argue, therefore, that there is a need to better understand and model the interaction of phylogeny with species’ traits and community composition.

**Innovation:** We outline a new method that identifies clades with unusual ecological structures, based around partitioning the variation of species’ site occupancies (*β*-diversity). Eco-phylogenetic theory would predict that these clades should also demonstrate distinct evolutionary trajectories. We suggest that modelling the evolution of independent trait data in these clades represents a strong test of whether there is an association between species’ ecological structure and evolutionary history.

**Main conclusions:** Using an empirical dataset of mammals from around the world, we identify two clades of rodents that tend not to co-occur (are phylogenetically overdispersed), and then find independent evidence of slower rates of body mass evolution in these clades. We suggest that our approach, which assumes nothing about the mode of species’ trait evolution but rather seeks to explain it using ecological information, presents a new way to examine eco-phylogenetic structure.

## 2 Introduction

Community phylogenetics (eco-phylogenetics) represents an attempt to link the evolutionary history of species to their present-day ecological interactions (Webb *et al*. 2002; Cavender-Bares *et al*. 2009). The field is young but controversial, and some of its fundamental assumptions have been criticised (notably by Mayfield & Levine 2010). Many community phylogenetic studies invoke niche conservatism (reviewed in Wiens *et al*. 2010) to assert that phylogenetic distance is a measure of distance in niche space, making phylogenetic structure a metric of ecological structure. However, few studies explicitly model such niche conservatism, and when a model is defined it is usually Brownian motion, which (arguably) describes neither niche conservatism (Losos 2008) nor similarity among distantly related species (emphasised by Godoy *et al*. 2014; Cadotte *et al*. 2017). Phylogeny is most often invoked as a proxy for unmeasured functional traits (as the ‘Phylogenetic Middleman’—Swenson (2013); see also Peres-Neto *et al*. (2012)). Such use undervalues phylogenetic relationships among species, which could be used to place (rather than approximate) species’ trait and distribution data within the context of past evolutionary and/or biogeographical processes that have shaped current patterns of species’ distributions and their co-occurrences. We cannot disentangle species’ functional trait evolution from their functional trait ecology if we use phylogeny as a measure of both. There is, therefore, a need to better integrate evolutionary history into community phylogenetics that parallels advances in the field of comparative analysis, where phylogeny is increasingly viewed as the inferential backbone for models of species’ trait evolution, not simply as a statistical correction (*e.g.*, Freckleton *et al*. 2011).

One of the earliest, and most commonly used, applications of community phylogenetic methods is to disentangle the impact of niche-based processes such as environmental filtering and competition on community assembly (Webb 2000; Cavender-Bares *et al*. 2006). Here, it is assumed that a community of closely-related species (phylogenetic clustering) reflects environmental filtering on the basis of phylogenetically conserved traits, while the converse (phylogenetic overdispersion) implies competitive exclusion (Webb *et al*. 2002). This direct mapping of phylogenetic structure onto ecological processes has been criticised (Cavender-Bares *et al*. 2009; Mayfield & Levine 2010), leading many to separately calculate the phylogenetic and functional trait structures of communities and then compare them (*e.g.*, Kraft & Ackerly 2010; Graham 2012). Critically, such comparisons do not capture the *interaction* between functional traits and phylogeny: how different ecological structures in different clades may have arisen. Multiple ecological and evolutionary processes interact to affect eco-phylogenetic structure, obscuring the signal of each process (Webb *et al*. 2002; Kraft *et al*. 2007; Cavender-Bares *et al*. 2009; Kembel 2009; Elliott *et al*. 2016). At its simplest, one clade may be functionally or phylogenetically overdispersed while another is clustered: only a clade-based approach can detect and unpick these conflicting signals. Figure 1 gives a conceptual example of how common ecological processes can produce variation among clades’ eco-phylogenetic structure. Using differences in ecological pattern among clades to guide questions about ecological assembly is a form of phylogenetic natural history (Uyeda *et al*. 2018).

**Figure 1.**
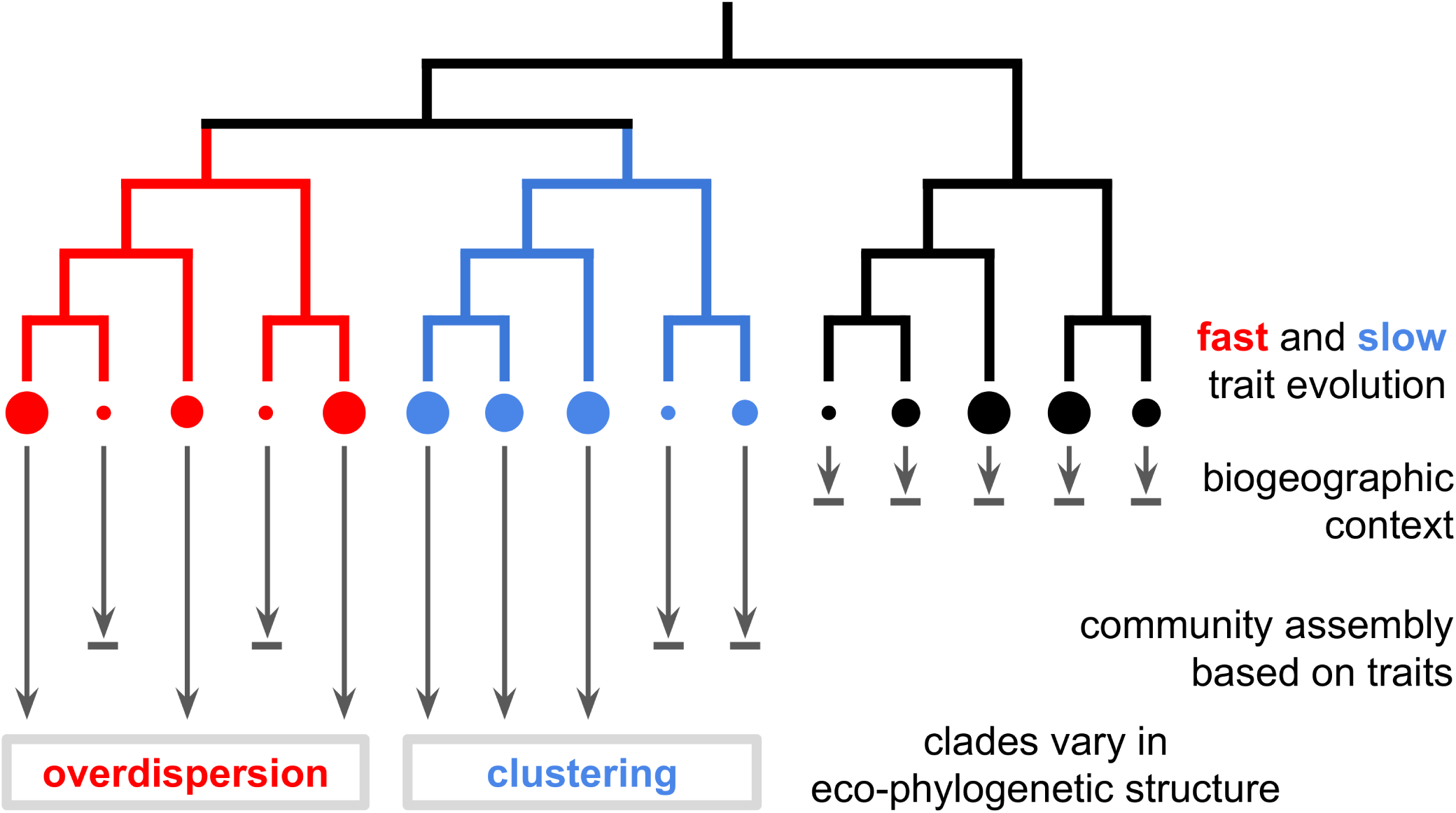
Linking clades’ evolution and ecological structure. Here we give an example of how clade-level variation in ecological structure (the tendency for close/distant relatives to co-occur) might arise. We consider a set of species that are initially filtered within some biogeographic (or meta-community) context; perhaps the clade is widespread but not all its members are present in every continent/region, for example. A trait, represented by the size of the circles at the tips of the phylogeny, evolves across the phylogeny, but evolves faster in one clade (the red branches) and slower in another (the blue branches). Ecological community assembly on the basis of this trait, regardless of mechanism, will result in different eco-phylogenetic structures across these clades. Re-framing our eco-phylogenetic analysis in terms of clades allows for the generation of falsifiable hypotheses about how species’ ecology and evolution interact. In this study, we use evidence of variation in the ecological structure of clades to test for variation in the evolution of those traits. It would also be possible to find clades with differing evolutionary patterns, and then use these to test for differing methods of ecological assembly and co-existence within those same clades. We emphasise that this diagram is but one example of how ecological assembly and the macro-evolution of species’ traits could interact. While we do not show the interaction of fitness and niche differences on species’ co-occurrence (*sensu* Chesson 2000; Mayfield & Levine 2010), we see no reason our approach could not be applied to more complex models of ecological assembly.

It is already well-appreciated in the eco-phylogenetic literature that different clades might demonstrate conflicting patterns, hinting at the interaction of ecological and phylogenetic structure (Ndiribe *et al*. 2013; Elliott *et al*. 2016). For example, the phylogenetic scale (*e.g.*, clades with different crown ages) of a study, and its relationship with spatial scale (*e.g.*, spatial extent) has itself become an object of study (see Swenson *et al*. 2006; Vamosi *et al*. 2009; Graham *et al*. 2018). Parra *et al*. (2010) were one of the first to examine the contribution of different clades to a single metric of phylogenetic structure. Later work expanded node-based analysis to consider the separate structures of individual clades (Pearse *et al*. 2013), and others have examined variation in environmental and biogeographic structure among clades (Leibold *et al*. 2010; Borregaard *et al*. 2014). Surprisingly, these advances in the measurement of clade-based eco-phylogenetic structure have been disconnected from clade-based advances in trait evolution (*e.g.*, Beaulieu *et al*. 2012; Mazel *et al*. 2016) and phylogenetic diversification (*e.g.*, Rabosky 2014). This is despite early work linking the order of trait evolution to community composition (Ackerly *et al*. 2006; Silvertown *et al*. 2006).

We suggest that one central assertion of community phylogenetics is that the evolution of species’ traits can be meaningfully linked to their present-day ecological structure (Webb *et al*. 2002; Cavender-Bares *et al*. 2009). A strong test of this assertion would be to link variation in the tempo or mode of trait evolution among clades with independent evidence of variation of community composition within those same clades. This goes beyond independently testing for phylogenetic structure of assemblages and traits (Swenson 2013): it tests hypotheses that specific clades should have different modes of trait evolution that cause, or are the consequence of, changes in the community composition of those clades (see figure 1). This approach looks to validate the assertion that variation among clades’ ecological structure is a product of the interaction of phylogeny with ecology using independent trait data.

In this paper, we extend an existing *β*-diversity framework (Legendre & De Cáceres 2013) to identify the unique contribution of phylogenetic clades to variation in community composition. Thus the contributions of clades can be compared with those of species and sites. Using this method it is possible to detect clades whose species do and do not tend to co-occur (‘clustered’ and ‘over-dispersed’ clades; Webb *et al*. 2002), and thus detect and disentangle variation in ecological structure across the tree of life. We apply our method to global mammal data (Jones *et al*. 2009; Thibault *et al*. 2011), where we find support for slower rates of body mass evolution in over-dispersed clades. By linking variation in clades’ ecological structure to variation in clades’ trait evolution, we show the power of phylogeny as data to help understand the evolution of ecological community assembly.

## 3 Methods

All software referred to below in *italics* are packages for the *R* environment (R Core Team 2017), and novel code written for this project is released in *pez* (in the function family *beta.part*; Pearse *et al*. 2015, to be added after acceptance, and currently in the supplementary materials). The supplementary materials contain code (that, using *suppdata*, also fetches all data; Pearse & Chamberlain 2018) that reproduces our empirical example in its entirety.

### 3.1 Overview and motivation

We often wish to determine whether species within an assemblage are more (phylogenetically *clustered*) or less (*overdispersed*) related compared to some expectation of assembly from a larger set of species, from which patterns we hope to infer some ecological mechanism. However, there is a growing understanding that such patterns are not necessarily uniform among the clades within a phylogeny (Leibold *et al*. 2010; Parra *et al*. 2010; Pearse *et al*. 2013; Borregaard *et al*. 2014). Indeed, phylogenetic clustering is an inherent property of *clades*: a phylogenetically clustered assemblage must have one or more over-represented clades, since clades are closely-related species and phylogenetic clustering is the presence of closely-related species. Below we describe how these patterns of clustering and overdispersion map onto clades within a phylogeny, using an extension of existing approaches to partition *β*-diversity (Legendre & De Cáceres 2013). By testing for differences in the evolution of such clades, we are able to test for associations between ecological and evolutionary processes, moving phylogeny from proxy for traits to data to be explored in the context of traits.

Figure 2 shows two assemblages (‘A’ and ‘B’) in an eight-species phylogeny; one of the clades is clustered, the other overdispersed. The general principle is clearer with species’ presence (‘1’) and absence (‘0’) data, but the calculations are the same with species’ abundances. While the variance (*σ*^2^) of each species’ occupancy of the two sites is the same (1/2), by summing the species’ occupancies within each clade the variance increases in the clustered clade and decreases in the overdispersed clade. When compared with simulations that provide null expectations of the expected variance in different clades, it is therefore possible to locate significant clustered and overdispersed clades across a set of ecological assemblages. We note that the standard advice when calculating *β*-diversity of abundance data is to work with a transformed data matrix (typically a Hellinger transformation; Legendre & Gallagher 2001). We do not do so here for clarity, and note that our simulations indicate our method is robust to such untransformed data.

**Figure 2.**
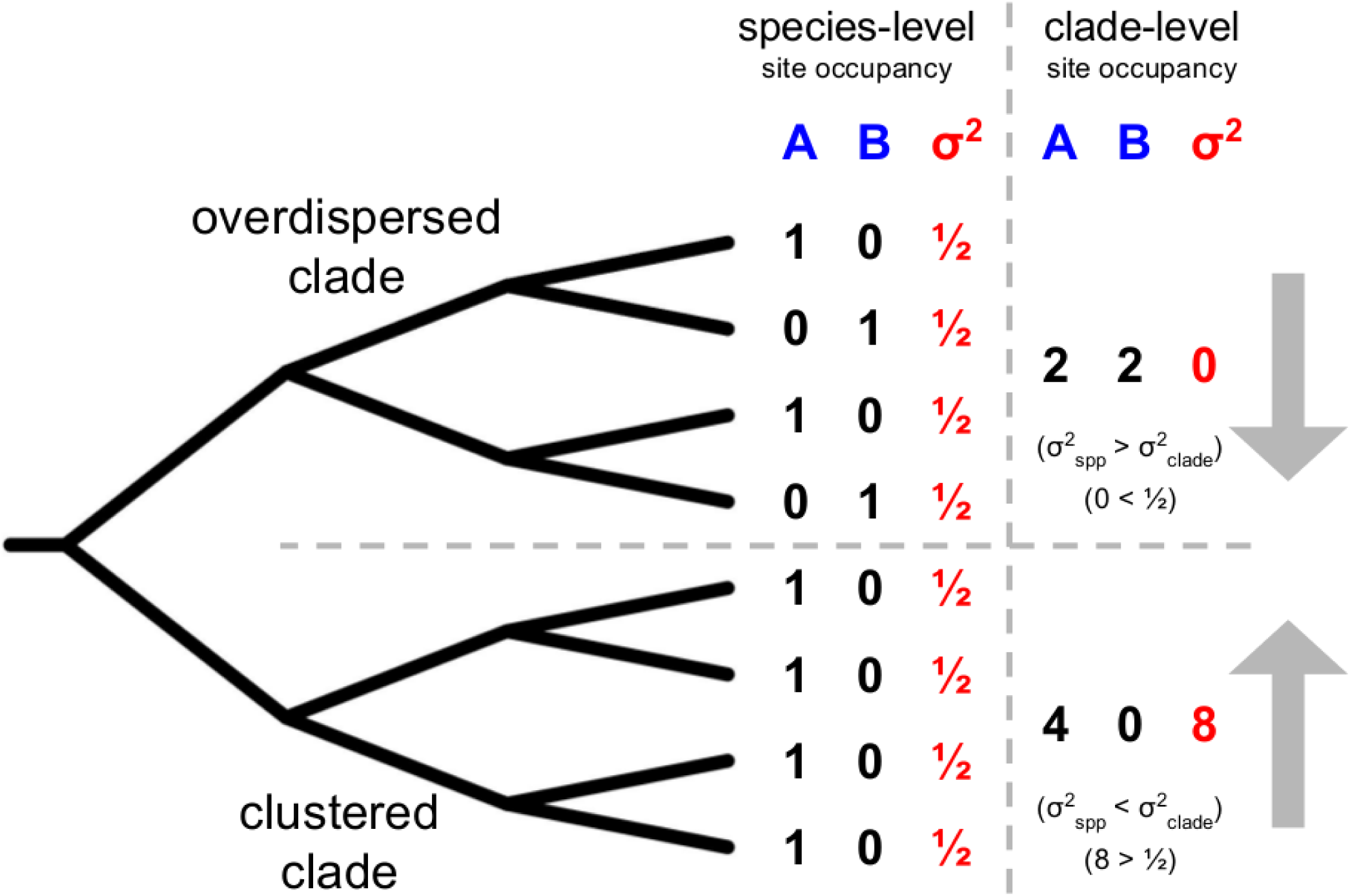
Overview of variance-based partitioning method. A horizontal dashed line splits the phylogeny into two clades: one has an overdispersed community phylogenetic structure (close relatives are unlikely to co-occur), and the other a clustered structure (closed relatives are likely to co-occur). It is these two kinds of ecological structure that our method aims to detect. A vertical grey dashed line separates species and grouped clade calculations. To the left of the vertical line, the abundances of each species in two assemblages (A and B) are shown alongside the variance (*σ*^2^) of each species’ compositions across the assemblages; all species have the same variance (½). To the right of the vertical line, community abundances for the species have been summed: the variance of these abundances is now much lower for the over-dispersed clade and much higher for the clustered clade. For simplicity, we use binary presence-absence data as an illustration, but this method can be applied to species’ abundances within assemblages. While there is an analytical expectation for clade-level variances (see text) we recommend using ecological null models to assess the significance of clade-level patterns. Note that when more than two sites are considered, a single variance value for each species is calculated across all species’ presences and absences (or abundances).

Once clades with different ecological structures have been identified, we can test whether the evolution of *independent* trait data differs within those clades (following Beaulieu *et al*. 2012). It is, of course, equally possible to test for variation in the evolution of clades first, and then to test the community composition of those clades using our *β*-diversity approach as the two procedures are performed independently. In such cases, clades with outliers in a PGLS regression (see Freckleton *et al*. 2011), or the output from methods such as SURFACE (Ingram & Mahler 2013), bayou (Uyeda & Harmon 2014), or `BAMM` (if shifts in speciation/extinction were of interest; Rabosky 2014) could be used to select candidate clades. Such clade-level tests directly map variation in ecological and evolutionary structure onto each other. Within this framework, phylogeny is not a mere proxy for missing species’ trait data (Mace *et al*. 2003; Srivastava *et al*. 2012; Swenson 2013): the interaction between phylogenetic, community composition, and trait data provides novel insight into how evolutionary history is linked with ongoing ecological processes.

We suggest that the main source of novelty in our approach is the comparison of trait evolution among clades with the ecological structure of clades. Additionally, our method of detecting variation among clades’ ecological structure is also novel. While there exist various approaches for identifying clades with particular ecological structures, our method is distinct from them. Firstly, and most importantly, it is a method for detecting variation in clade-level compositions (*c.f.* Ives & Helmus 2011). Secondly, it compares multiple sites (*c.f.* Pearse *et al*. 2013) simultaneously as it measures *β*-diversity. Thirdly, it does not seek to find clades that contribute to an overall pattern (*c.g.* Parra *et al*. 2010) but rather identify contrasting patterns among clades. Finally, it models all species simultaneously and so does not compare species’ individual drivers of presence/abundance, making it capable of detecting overdispersion (*c.f.* Leibold *et al*. 2010; Borregaard *et al*. 2014).

Because our clade-wise test of ecological structure is novel, so too are our definitions of overdispersion and clustering (*c.f.* Webb 2000; Webb *et al*. 2002; Cavender-Bares *et al*. 2009). Here we define a clustered clade not on the sole basis of presences within a single site, but rather the pattern of presences and *absences* across *multiple* sites. For example, the clustered clade in figure 2 would not traditionally have been considered clustered in site B. Further, we emphasise that, in our framework every clade has a variance, and while we present “clustered” and “overdispersed” clades in figure 2, it is likely that many clades fall somewhere between these two extremes.

### 3.2 Extensions of *β*-diversity and significance tests

The method of Legendre & De Cáceres (2013) is essentially based around variance in species’ abundances across sites. In this context, our *β*-diversity partitioning extends species-level contributions to consider clade contributions. This allows ecologists interested in comparing the contributions of species and sites to *β*-diversity patterns to also compare the contributions of clades. Indeed, while we focus solely on phylogenetic clades in this manuscript, we see no reason why this approach could not be applied to other (hierarchical) groups of species, such as those produced using functional traits (Petchey & Gaston 2006).

We suggest two ways to assess the significance of a clade’s departure from the expected variance (the clade-level variances, *σ*^2^, on figure 2). The first is an ‘exact’ method based on the expectation of variances, and is described in the Supplementary Materials. The second method is based on the contrast of observed clade variances with null distributions of variances estimated via permutation (*e.g.*, reshuffling species’ identities across the phylogeny, reviewed in Gotelli 2000). Ranking a clade’s observed variance among its null variances would reveal whether a clade had unusually high or low variance. The null model approach protects against cases in which a clade whose members are entirely absent or omnipresent within a set of communities from being highlighted as a clade with low variance. We strongly recommend the use of ecological null models to assess significance as they allow more flexibility over the processes contained with a null hypothesis, and are not as reliant on the statistical distribution of species’ abundances.

### 3.3 Simulations testing clade-partitioning

We used simulations to verify our method’s ability to detect variation in ecological structure among clades. Below we describe each parameter of the simulation, listing each parameter in *italics* and its values across the simulations (in parentheses). We simulated phylogenies of *n*_*spp*_ species (either 50 or 100) following a pure-birth Yule process (using *geiger v. 2*; Pennell *et al*. 2014). We then selected a focal clade containing either 5–10% or 10–20% of the species in the phylogeny, and simulated a trait under Brownian motion (root set to 0, also using geiger v. 2; Pennell *et al*.2014) across the entire phylogeny with a *σ*^2^ 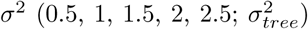excluding the focal clade, which had a *σ*^2^ a multiple of 10 greater or lesser than across the entire tree 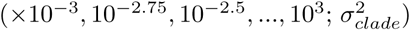We then simulated community assembly across *n*_*site*_ sites (either 50 or 100) on the basis of this simulated trait: in each site we randomly selected a species and randomly drew species to be members of that community on the basis of their trait distance from that species. Thus species with absolute differences in simulated traits *≥* 1 from the focal species would have a probability of membership of 0; a species with a difference of |0.5| would have a probability of 0.5. We acknowledge that this mapping between trait difference and probability of co-occurrence is arbitrary and was chosen for the sake of simplicity. In related simulations, however, we have seen little evidence that this qualitatively affects our method’s performance.

These simulations represent a form of ecological assembly that is deliberately agnostic with regard to any particular ecological mechanism (*e.g.*, facilitation, competition, or environmental filtering), but conceptually similar to that shown in figure 1. It is well-known that multiple ecological mechanisms can result in the same eco-phylogenetic structure (Cavender-Bares *et al*. 2009; Mayfield & Levine 2010). Here we simply aim to simulate variation and then test our power to detect that variation, given pattern detection is an important aid in determining the processes underlying community assembly. A clade can evolve faster than the rest of the phylogeny 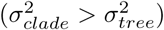, in which case we would expect close-relatives to rarely co-occur within a clade (an overdispersed clade; see figure 2). A clade can also evolve slower than the rest of the phylogeny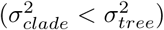, in which case we would expect close-relatives to frequently co-occur (a clustered clade; see figure 2). Even in simulations where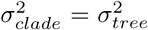, we still evolved a separate trait for the focal clade, making this an extremely conservative test of our method.

We repeated this simulation approach for all combinations of our parameter values, and an additional 20 times for each combination with identical 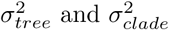, resulting in a total of 2160 simulations. For each simulation, we ranked the observed variance of the focal clade within 99 randomisations (the observed value was included as part of the null distribution, totalizing 100 values for each null distribution), swapping species’ identities on the phylogeny and keeping everything else constant. These rankings provide probabilities under the null hypothesis: values greater than 0.975 suggest clustering (at *α*_5%_) and values lesser than 0.025 suggest overdispersion. This provides a test of whether our method can reliably detect overdispersion (ranked the lowest or second-lowest variance in the randomisations when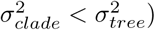 clustering (ranked the highest or second-highest variance in the randomisations when 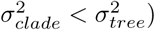 and whether it is vulnerable to false-positives (*i.e.*, type I error rates greater than the expected confidence level at 0.05—ranking consistent with clustering or overdispersion when 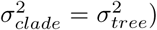. Note that clades are hierarchically nested within each other, and so species that are shared across clades mean clades’ structure are not necessarily independent. While we make reference to this in the discussion, we do not conduct simulations to investigate this further, as it is a feature that has been discussed at length in the literature (*e.g.*, Alfaro *et al*. 2009). We draw the reader’s attention to the fact that we conduct these simulations over a range of parameter values, with the explicit aim of finding the conditions under which our method performs well and where it underperforms.

### 3.4 Empirical example: global rodent communities

To provide an empirical example of how our approach uses ecological structure to generate specific hypotheses about the evolution of species’ traits, we present an analysis of a global rodent dataset. We took data from a global mammal community dataset (Thibault *et al*. 2011), global phylogeny (Bininda-Emonds *et al*. (2007), updated by Fritz *et al*. (2009)), and body mass from a large database for mammal traits (Jones *et al*. 2009). This global community dataset covers a number of continents and community types, and body mass is known to be a good proxy for ecological interactions in rodents (see Thibault *et al*. 2011). Excluding all species not covered in all three datasets left us with abundance information for 483 species across 939 sites (assemblages) worldwide. Following the method described above, we identified clades’ *β*-diversity and assessed statistical significance by comparison to 1000 independent-swap randomisations (Gotelli 2000; Kembel *et al*. 2010).

Identifying clades with unusual *β*-diversity does not, however, test whether there is an association between the evolution of a clade and its ecological structure. We therefore fitted Brownian motion and Ornstein-Uhlenbeck (OU) models using *OUwie* (Beaulieu *et al*. 2012) to the (log-transformed) body mass data, contrasting models with shared and varying parameters for clades identified as significantly departing from our null expectations in our *β*-diversity approach. Support for Brownian and OU models with different parameters for the clades identified by the *β*-diversity ecological analysis would suggest a link between ecological structure and trait evolution. Where hierarchically-nested clades were identified, we selected the oldest clade as this is more conservative (the ‘cascade’ problem; see Discussion) and parameter estimation is more accurate in larger clades (Beaulieu *et al*. 2012). In the Supplementary Materials, we present results of a series of permutation tests we performed to ensure that our evolutionary model-fitting was not biased towards finding support for particular evolutionary hypotheses.

## 4 Results

Results from our simulations are presented in table 1 and figure 3, and show that our method powerfully and reliably detects variation in phylogenetic structure among clades. Our method has strong statistical power to detect clustering (higher variance within a clade; the red line in figure 3), and a somewhat reduced power to detect overdispersion (lower variance within a clade; the blue line in 3). As shown in table 1, however, greater sampling modifies this: sampling 100 species across 100 sites additively increases the ranking of the observed variance by 10% (*e.g.*, from *p* = 0.85 to *p* = 0.95) in comparison with 50 species across 50 sites. Our method shows a tendency to spuriously suggest support for clustering (*i.e.*, overall inflated type I error rates in simulations of 24% at two-tailed *α*_5%_; see figure 3), but again this varies depending on the biological context. As shown in table 1, focal clades that make up large proportions of the total data are more likely to be erroneously identified as clustered: if the focal clade contains 10 of the 100 species in a system (*n*_*sites*_ = 50, *σ*^2^=1) the predicted probability of clustering is 0.77, but if the clade contains 20 species (*i.e.*, 20% of the species) that prediction rises to 0.95. Neither of these predicted values are statistically significant at *α*_5%_. As we highlighted above, we conducted simulations across a wide region of parameter space to highlight where our method performs well and where it performs poorly. Thus the raw results plotted in figure 3 do not necessarily reflect our average expectations for performance of our method.

**Table 1:** Simulations showing how method performance varies as a function of phylogeny and clade size, rate of trait evolution, and effect size. Each sub-table shows the results of modelling the estimated probabilities that focal clades are clustered (higher variance; a), overdispersed (lower variance; b), and random (null, no difference; c) across the simulations. At an *α*_5%_, a predicted probability of 0.025 or 0.975 would provide statistical support for the focal clade being clustered or overdispersed, respectively. Generalised Linear Models with a quasibinomial error structure were used to account for non-normality of errors in the clustering (a) and overdispersion (b) models, and so coefficients are reported on the logit scale. In (a), a greater statistical power to detect clustering is most strongly associated with the number of species in the focal clade and the difference in evolutionary rate between the focal clade and the rest of the phylogeny (deviance: *null*_527_ = 98.46 and *residual*_522_ = 62.52; estimated *dispersion* = 0.51). In (b), a greater statistical power to detect overdispersion is most strongly associated with the difference in evolutionary rate between the focal clade and the rest of the phylogeny and the number of sites sampled (deviance: *null*_524_ = 277.74 and *residual*_519_ = 152.97; estimated *dispersion* = 0.51). In (c), there is a slight tendency for larger focal clades to appear more clustered, and for faster-evolving traits to drive overdispersion, even when focal clades evolve under the same model as the rest of the phylogeny (*F*_4,919_ = 13.75; *r*^2^ = 5.64%; *p <* 0.0001). We recommend that more attention should be paid to effect estimates than statistical significance in these models, since statistical significance can be driven by sample size and these are the results of simulations.

| | Estimate | Std Err | $z$ | $p$ |
| --- | --- | --- | --- | --- |
| Intercept ($n_{spp} = n_{sites} = 50$) | -0.0574 | 0.5643 | -0.10 | 0.9191 |
| $\log_{10}(\frac{\sigma_{tree}^2}{\sigma_{clade}^2})$ | 0.8788 | 0.2015 | 4.36 | <0.0001*** |
| $n_{clade}$ | 0.3710 | 0.0916 | 4.05 | 0.0001*** |
| $\sigma_{tree}^2$ | -0.1703 | 0.2753 | -0.62 | 0.5363 |
| Contrast—$n_{spp} = 100$ | 0.3772 | 0.5263 | 0.72 | 0.4739 |
| Contrast—$n_{sites} = 100$ | 0.4182 | 0.3122 | 1.34 | 0.1809 |

(a) Clustering (higher variance)
| | Estimate | Std Err | $z$ | $p$ |
| --- | --- | --- | --- | --- |
| Intercept ($n_{spp} = n_{sites} = 50$) | 1.2188 | 0.2701 | 4.51 | <0.0001*** |
| $\log_{10}(\frac{\sigma_{clade}^2}{\sigma_{tree}^2})$ | -1.8568 | 0.1265 | -14.68 | <0.0001*** |
| $n_{clade}$ | 0.0048 | 0.0244 | 0.20 | 0.8437 |
| $\sigma_{tree}^2$ | -0.1751 | 0.1399 | -1.25 | 0.2114 |
| Contrast—$n_{spp} = 100$ | -0.1595 | 0.2007 | -0.79 | 0.4271 |
| Contrast—$n_{sites} = 100$ | -0.3402 | 0.1569 | -2.17 | 0.0306* |

(b) Overdispersion (lower variance)
| | Estimate | Std Err | $z$ | $p$ |
| --- | --- | --- | --- | --- |
| Intercept ($n_{spp} = n_{sites} = 50$) | 0.6432 | 0.0296 | 21.72 | <0.0001*** |
| $n_{clade}$ | 0.0174 | 0.0030 | 5.90 | <0.0001*** |
| $\sigma_{tree}^2$ | -0.0338 | 0.0170 | -1.98 | 0.0478* |
| Contrast—$n_{spp} = 100$ | -0.0088 | 0.0239 | -0.37 | 0.7123 |
| Contrast—$n_{sites} = 100$ | 0.0222 | 0.0191 | 1.16 | 0.2455 |

**Figure 3.**
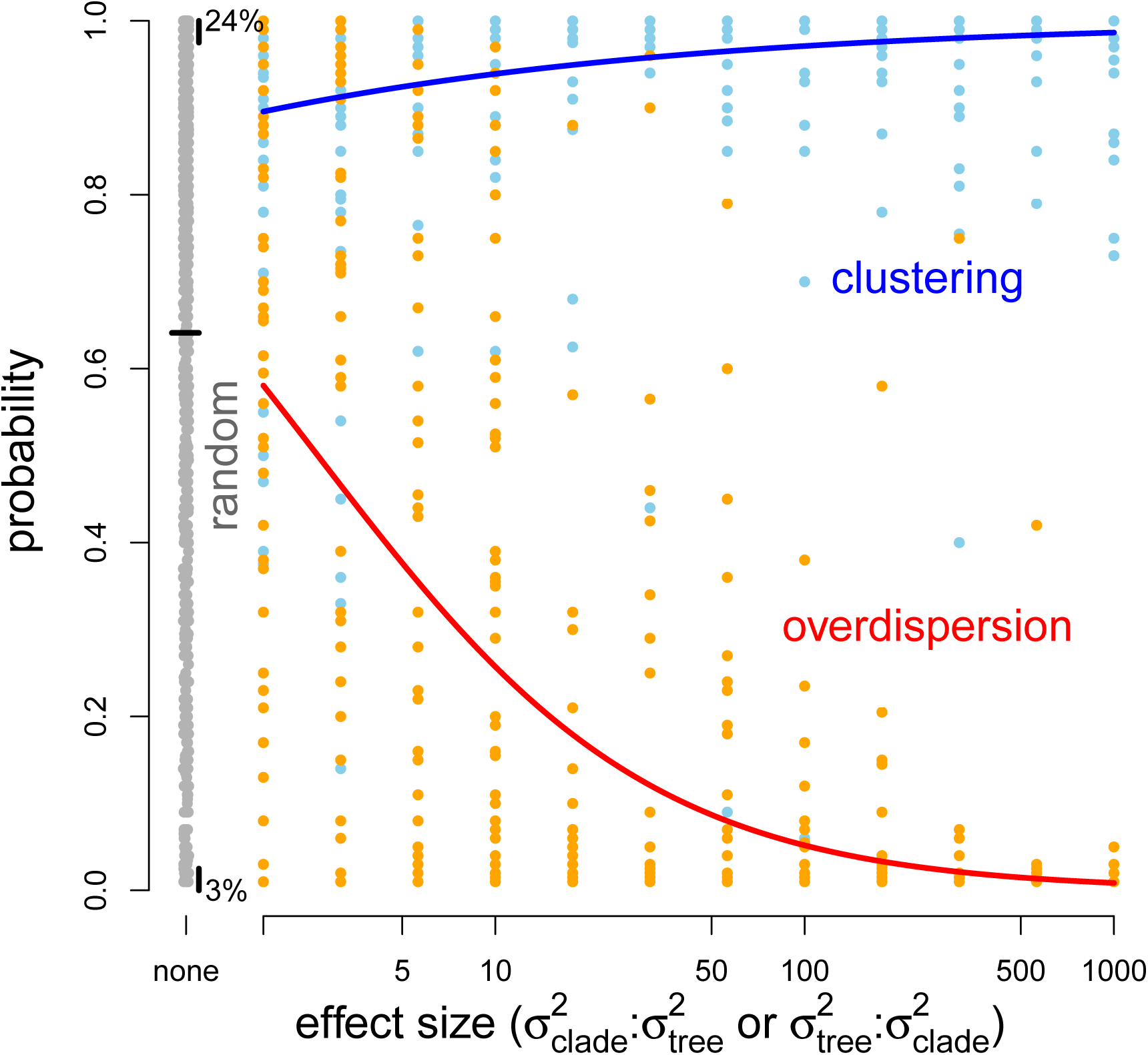
Simulations showing how method performance increases with effect size. In grey, the probabilities are shown for when there was no difference between the model of trait evolution in the focal clade and the rest of the phylogeny. The mean of these values, along with the percentage of values lying beyond the 2.5% and 97.5% quantiles, are shown in black. In blue, the probabilities for the overdispersed 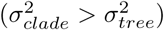 are shown, along with a quasi-Binomial GLM prediction. In red, the probabilities for the clustered (high variance;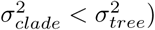 are shown, along with a quasi-Binomial GLM prediction. At an *α*_5%_, a predicted probability of 0.025 or 0.975 would provide statistical support for the focal clade being clustered or overdispersed, respectively. None of these curves account for the additional explanatory variables used in the models in table 1, and thus these curves are conservative. Raw data used to parameterise the models shown in table 1.

From our analyses of the rodent dataset, figure 4 shows the mammal clades identified as having significantly different variance distributions, and table 2 shows the AIC comparison of models of trait evolution that incorporate variation within those clades. The two clades on which we focused (marked on figure 4) are the *Scuridae* (squirrels) and their sister family the *Gliridae* (dormice), and the *Echimyidae* (a Neotropical rodent family) and some close-relatives within what is sometimes called the *Caviomorpha* (*e.g.*, South American rodents like the guinea pig). We refer to these two groups as the ‘squirrels’ and ‘cavis’, respectively, although these terms do not precisely map onto all species within the clades. The squirrel and the cavi clades were both identified as having low variance (phylogenetic overdispersion). Note that our method also detected clades indicative of clustering (high variance). As the low-variance clades are nested within these high-variance clades, we suggest they might reflect important eco-evolutionary shifts. The detection of both phylogenetic clustering and overdispersion demonstrates the ability of our method to reveal both kinds of pattern in empirical datasets.

**Table 2:**
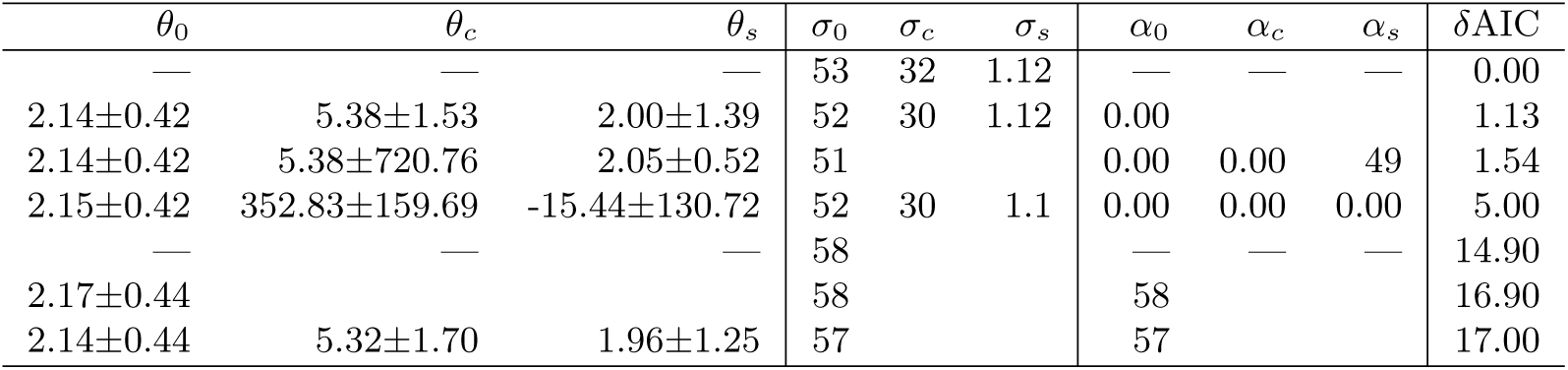
Results of log(body mass) evolutionary modelling. Above are the *θ* (optimum), *σ* (rate), and *α* (rate of return to optimum) estimates, along with AIC and *δ*AIC values, for all trait evolutionary models. Each row represents a different model; ‘—’ is used to indicate when a parameter is not fit in a model, and where only a single estimate for a parameter is given (*e.g., θ*_0_) only a single parameter was fit across the whole phylogeny. Thus rows one and four represent Brownian motion (models with no optima), and all other rows are variants of Ornstein-Uhlenbeck models. In subscripts of parameters, ‘c’ refers to the ‘capi’ clade, ‘s’ to the ‘squirrel’ clade, and ‘0’ to the remainder of the phylogeny. See text and figure 4 for a description of these species making up each clade. The *α* and *σ* estimates have been multiplied by 10^−4^ for brevity of presentation. The four most likely models according to *δ*AIC all contain clade-level variation, strongly supporting different patterns of evolution in the clades highlighted by the clade-level partitioning of *β*-diversity(see text).

**Figure 4.**
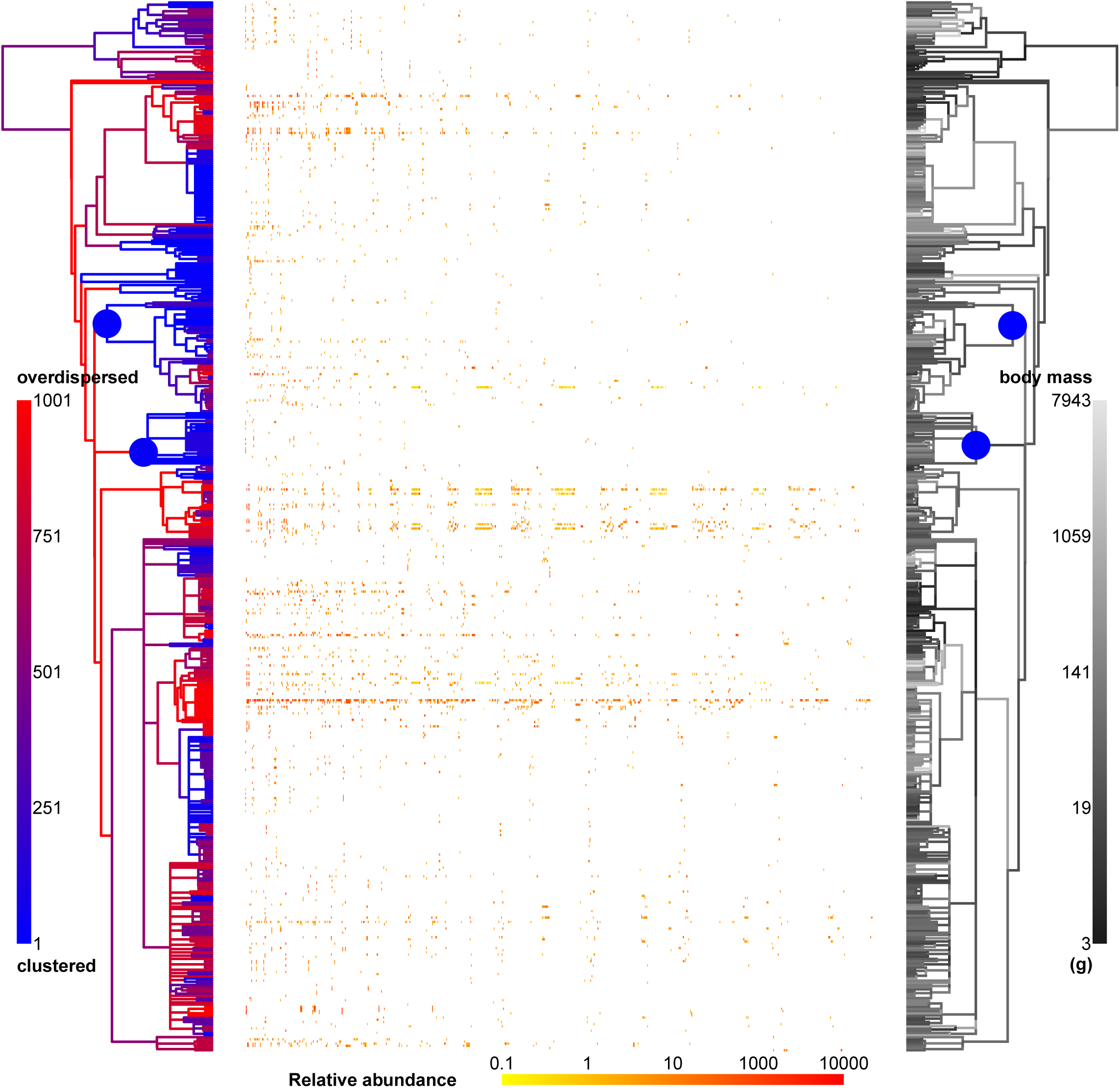
Empirical mammal results showing associations between ecological structure of clades and their rates of body mass evolution. To the left and right, the phylogeny of all 483 mammals in the study. Two large blue circles on the nodes of each phylogeny indicate the two ‘squirrel’ and ‘cavi’ clades tested in the evolutionary analysis (see text and table 2). The left-hand phylogeny is coloured according to the ranking of the clades’ variances; a ranking of 1 (blue; see legend) would indicate a clade whose variance was lower than all 1000 null permutations, and a ranking of 1001 (red; see legend) a clade whose variance was higher than all 1000 null permutations. In the center, a site-by-species matrix of relative abundance in all 939 assemblages, with a colour-scale indicating relative abundace (see legend at bottom). The right-hand phylogeny is shaded according to a reconstruction of body mass (g) across the phylogeny (using phytools Revell 2012). Although this reconstruction does not explicitly model variation in rate among clades, variation in size across its branches can be seen.

Table 2 shows that the squirrel and cavis clades were also characterised by different rates of trait evolution. The top four models, with *δAIC* less than 5, all supported different rates of body mass evolution for these two clades. The fifth-best model had a *δAIC* of 14.9 and so there is only limited support (Burnham & Anderson 2002) for the alternative hypothesis that trait evolution is constant across the squirrels, cavis, and the rest of the mammal phylogeny. The lowest-AIC model favoured a simple three-rate Brownian motion model in which the rate of body mass evolution in squirrel and cavi clades is significantly slower, most notably in the squirrel clade. Our permutations procedure suggest that, by chance, we would expect to find the *opposite* pattern to that found with the empirical data (see Supplemental Materials), giving greater strength to our findings.

## 5 Discussion

We have presented a novel method for identifying clades (groups) of species whose ecological community structure differs from other species across a set of communities. Simulating species’ phylogenies and responses to an environmental gradient, we demonstrated that the method reliably detects shifts in the variance of species’ abundances, identifying different phylogenetic structures. Most importantly, however, we have also shown, using empirical data, that the tempo of trait evolution shifts within clades associated with unusual present-day ecological structure. To the best of our knowledge, this is the first eco-phylogenetic framework that performs hypothesis tests of associations between the evolution and ecological community composition of clades. By linking variation among clades’ ecological structure with independent evidence for variation in those clades’ rates of trait evolution, we have found evidence for an interaction between evolutionary and ecological information. We argue that our approach, combining evidence of both ecological and evolutionary patterns, has more power to answer questions about the underlying eco-evolutionary drivers of community assembly than methods focusing singularly on phylogenetic or trait data alone.

### 5.1 β-diversity partitioning in community phylogenetics

The use of phylogeny as a proxy for ecological process has been criticised. It has been argued that there is little need for phylogeny if we already have functional traits (Swenson 2013), and phylogenetic pattern rarely maps directly onto ecological process (a critique that applies equally to functional traits; Mayfield & Levine 2010). However, we have suggested one central premise of community phylogenetics is that there is an association between the evolution of species’ traits and the phylogenetic structure of the communities in which they are found. Many community phylogenetic studies, like ours, examine the tempo and mode of trait evolution within their system (*e.g.*, Swenson *et al*. 2006; Kraft *et al*. 2007), but few have asked how trait evolution and community phylogenetic structure are linked and feedback into each other. Simple measures of phylogenetic signal assume complete, or at least unbiased, taxon sampling (Pagel 1999; Blomberg *et al*. 2003), and so eco-phylogenetic structure, which, by definition, implies nonrandom taxonomic representation, may mask broader (true) patterns of trait evolution by introducing non-random taxonomic sampling. Our approach offers a coherent framework to test for links between the macro-evolutionary dynamics of clades and their present-day (or relatively recent past) community composition.

Despite conceptual issues, the utility of phylogeny in predicting species’ traits (Guénard *et al*. 2013), Janzen-Connell effects (Gilbert & Webb 2007), invasion success (Strauss *et al*. 2006), and ecosystem function (Cadotte *et al*. 2013) suggests phylogeny will remain a useful (Tucker *et al*. 2018), if imperfect (Cadotte *et al*. 2017), proxy in ecology for some time. Yet we suggest that phylogeny is more than just a surrogate for unmeasured traits, and that it provides us with the ability to link patterns and processes in ecology and evolution. Here, we map patterns in separate ecological assemblages and species trait datasets onto each other, linking them by treating phylogeny in and of itself as data in two separate analyses. Our approach does not invoke niche conservatism, but rather seeks to understand how traits have evolved and match with (current) patterns of species co-occurrences across local communities. As such, there is no requirement that closely related species are more ecologically similar or compete more strongly, assumptions that have been heavily criticised (Cahill *et al*. 2008; Mayfield & Levine 2010). Our results simply support a link between the ecological structure (*β*-diversity) of clades and the evolutionary history of those clades. The evolutionary patterns we observe come from interactions, or the absence of interactions, that occurred over millions of years, potentially in assemblages very different to those we see today. Our analyses indicate that these past interactions have left an imprint on present-day ecological structure, and imply that future evolutionary trajectories may be influenced by present-day species interactions.

In our analysis of small mammal assemblages, we showed that the ‘capi’ and ‘squirrel’ clades, whose members tended not to co-exist (their clade variances were low), have lower rates of character evolution (table 2). Rodent body size is a driver of ecological competition (Bowers & Brown 1982; Ernest 2005), and our results suggest slower evolution of body size is a driver of variation in the present-day community composition of our small-mammal communities. The clades we have focused on are relatively small and young (see figure 4), and previous work (Ackerly *et al*. 2006; Silvertown *et al*. 2006) has suggested that traits that evolve early and late in the evolutionary history of a clade may affect ecological assembly differently. Our results imply that it is not just the timing of body size evolution that may be important, but also its rate of evolution. We do not yet know what caused this slow-down in the ‘capi’ and ‘squirrel’ clades and whether these associations are driven by changes in diversification rate (which can be confounded with trait evolution; FitzJohn 2010). There is, however, some evidence that younger clades tend to co-occur more than older ones (Pearse *et al*. 2013; Parmentier *et al*. 2014).

### 5.2 Method perfprmance

We show that our method has good statistical power, and compares favourably to the widely used NRI and NTI metrics of phylogenetic community structure, for which statistical power has been estimated at less than or equal to 20% (Kraft *et al*. 2007) and 60% (Kembel 2009). In some cases, however, we observed inflated type I error rates relative to these other methods (see below for discussion). In many ways these are unfair comparisons, given that our approach makes use of information from multiple sites (although the number of species with phylogenetic structure is comparable), which we would argue this is a strength of our method. Phylogenetic Generalised Linear Mixed Models (Ives & Helmus 2011) also use many sites at once, and our results compare favourably to this approach (87% detection rate for phylogenetic clustering, 53% for overdispersion, but with fewer sites than in our study). It is important to note, however, that these alternative methods are intended to answer different questions. We make these comparisons simply to demonstrate that our approach performs reasonably in comparison with others, even in simulations where the number of species in a focal clade could be as low as 5 and the datasets themselves are small (no more than 100 species or sites).

Our simulations show that, in cases where the focal clade makes up a large proportion of the species under study (in our simulations, over 20%) type I error rates could be inflated. We do not feel that this is of concern, for several reasons. First, within our framework, clades must be detected as significant both in terms of their ecological structure and also their historic trait evolution. As such spurious identification of structured clades would tend to weaken any association between their ecology and evolution. Second, it is rare that ecological assemblages are truly randomly structured: the norm is for them to display some degree of phylogenetic structure (Vamosi *et al*. 2009). We suggest most biologists may be more interested in detecting the difference between overdispersion and clustering, not overdispersion/clustering versus random assembly. This is the case in our empirical example, where we examined clades that were overdispersed whose sisters are clustered. Third, we used two separately evolved traits for the tree and the focal clade. Our method may be detecting genuine differences between the focal clade and the tree as a result of different ecological assembly on the basis of genuinely different traits. We also note that type II error rates have been shown to be even higher for other, more commonly used, metrics of phylogenetic structure. For example, *SES*_*MP* *D*_ and *SES*_*MNT* *D*_, when estimated by taxa-shuffling null distributions such as we employ here, can have type I error rates of c. 50% (Kembel 2009).

### 5.3 Potential methodological extensions

Like similar approaches (Parra *et al*. 2010; Pearse *et al*. 2013; Borregaard *et al*. 2014), does not directly consider nestedness (Ulrich *et al*. 2009), where the significance of a clade ‘cascades’ up into higher super-sets of hierarchical groupings (*c.f.* the ‘trickle-down’ problem in diversification analysis; Purvis *et al*. 1995; Moore *et al*. 2004). One possible extension would be to compare each clade with the *summed* clades subtending it (not, as in the method we present, the species within it). Thus each clade in a fully resolved phylogeny would have its variance compared with the variances of the two clades subtending it (our supplementary code permits this). Significance could be tested through null permutation, as we use in this manuscript, or potentially through nested ANOVAs. However, we suggest that this cascading is not so much a limitation but rather a matter of interpretation; that a group has unusual *β*-diversity because it contains other unusual groups does not strike us as problematic. A balanced approach could limit the study to particular clades on the basis of age or other *a priori* interest, or to hold problematic clades constant in null randomisations.

We also note that our approach for identifying ecologically structured clades does not incorporate phylogenetic branch lengths. Branch lengths inform models of trait evolution, and so for our purposes of mapping *independent* evolutionary structure onto ecological structure we consider it undesirable to have branch lengths play a role in both aspects. For those interested in incorporating branch lengths in other situations, a simple approach would be to multiply each species’ abundance by its evolutionary distinctiveness (Isaac *et al*. 2007) or another measure of its phylogenetic uniqueness (*e.g.*, Redding & Mooers 2006; Cadotte *et al*. 2010). However, depending on the question at hand this might ‘average out’ the signal that we may be interested in detecting. For example, if community composition varies with phylogenetic scale (Webb *et al*. 2002; Cavender-Bares *et al*. 2009; Vamosi *et al*. 2009), it might be better to model the standard effect size (SES; *sensu* Kembel 2009) of node variance as a function of node age (see Pearse *et al*. 2013).

## 5.4 Conclusion

We suggest that the identification of clades with unusual ecological structure is of at least as much interest as the summary statistics that have been used frequently to describe phylogenetic assemblage structure but which map only poorly to ecological process. Further, we see that establishing links between assemblage structure and the evolution of species’ traits as a central premise of community phylogenetics, but has been rarely tested. As a field, community phylogenetics is well-placed to take advantage of recent advances in trait evolution (Pennell & Harmon 2013; Nuismer & Harmon 2015) and eco-phylogenetic theory (Pigot & Etienne 2015). We have outlined here an approach to directly test links between the ecological structure of assemblages and the evolution of species’ traits. As we gain a firmer grasp of assemblages’ phylogenetic structure, we can begin to model it as data, not merely measure its pattern.

## Data accessibility statement

No new data is released as part of this manuscript. All simulations and analysis R code is released in the supplement.

## Acknowledgements

We are grateful to XXX anonymous reviewers, and the editorial board, for their help improving this manuscript. TJD and the Davies lab are funded by Fonds de Recherche Nature et Technologies grant number 168004 and an NSERC Discovery Grant. WDP and the Pearse lab are funded by NSF ABI-1759965, NSF EF-1802605, and USDA Forest Service agreement 18-CS-11046000-041.

